# Distinct Effects of Aducanumab and Lecanemab on Intraneuronal Endogenous Aβ42 and Phosphorylated Tau in Alzheimer’s Disease Treatment

**DOI:** 10.1101/2024.02.06.579113

**Authors:** Ming-Jie Liu, Zhu Long, Fuyun Li, Xiaobang Shi, Yan-Fei Shen, Meng-Lu Liu

**Author notes:** Co-corresponding authors: Ming-Jie Liu, and Meng-Lu Liu, Address: 4/F Building 5, Chuang e Huigu, 777 Zhongguan West Road, Zhenhai District, Ningbo 315201, China. These authors contributed equally (Ming-Jie Liu, Zhu Long).

## Abstract

In the treatment of Alzheimer’s Disease (AD), two FDA-approved monoclonal antibodies, Aducanumab (Adu) and Lecanemab (LCN), exhibit significant differences in clinical benefits. By utilizing human-induced basal forebrain cholinergic neurons (BFCNs) derived from AD patient skin fibroblasts, we successfully recapitulated the natural endogenous neuropathologies of Aβ42 and Tau within just 21 days and revealed distinct intraneuronal effects of Adu and LCN. Both antibodies are internalized into BFCNs and localize with cytosolic Aβ42. However, LCN, selectively targeting Aβ42 oligomers and protofibrils, triggers TRIM21 pathway and significantly enhances autolysosome- and proteasome-mediated Aβ42 clearance, thereby leading to a marked reduction in phosphorylated Tau181 (pTau181) pathology. In contrast, the fibrillized Aβ42-selective Adu shows considerably weaker effects. This study not only reveals the unique intraneuronal actions of Adu and LCN but also provides a reliable and accessible human neuronal model for evaluating potential AD therapeutics, emphasizing the importance of intraneuronal pathology in the treatment of AD.

## Introduction

Alzheimer’s disease (AD) is characterized by the extracellular deposits of amyloid plaques and intracellular tau aggregates (neurofibrillary tangles) in the brain^1–4^. These hallmark pathologies affect millions worldwide, driving extensive research efforts to develop effective treatments^3–5^. Despite considerable advancements, there is still a substantial gap in our understanding of the intraneuronal effects of specific therapeutic interventions^3^, primarily attributed to the lack of neuron models that accurately replicate the natural endogenous intraneuronal pathology^6–8^. This gap is particularly evident in understanding how the FDA-approved monoclonal antibody drugs impact the pathology within neurons in AD.

Aducanumab (Adu), the pioneer FDA-approved monoclonal antibody for AD, targets amyloid plaques^9,10^. While it effectively diminishes the extracellular fibrillar Aβ42 load in the brain, its impact on clinical symptoms remains limited^9^. In contrast, Lecanemab (LCN) is a subsequent FDA-approved monoclonal antibody, initially formulated to target soluble Aβ42 protofibrils^11–14^. Uniquely designed as a bispecific antibody, LCN also enhances blood-brain barrier (BBB) permeability by recognizing the transferrin receptor^14^. Extensive research has shown that LCN possesses a strong affinity for oligomeric Aβ42^15^. It is distinguished by its dual ability to reduce brain Aβ42 protofibrils and inhibit the proliferation of Aβ42 fibrils^16^, showcasing superior clinical symptom improvement compared to Adu^11,17^. Other AD monoclonal antibody drugs, either in development or those that have failed in trials, typically focus on extracellular Aβ42 oligomers and protofibrils, or insoluble fibrillar Aβ42^13,18,19^. Growing research underlines the predominance of neuronal Aβ42 as a major contributor to brain amyloid^2,6,20–24^. Present in various forms such as soluble monomers, oligomers, and protofibrils, intraneuronal Aβ42 accumulation and aggregation precede the formation of amyloid plaques and are closely linked to early memory and cognitive decline in AD^2,6,20,21^. Amyloid plaques primarily derive from the residual Aβ42 following abnormal neuronal secretion or neuronal death^20–22^. This underscores the necessity for AD drug development to also address the accumulation and aggregation of intraneuronal Aβ42.

The development of disease-specific neuronal models that accurately replicate intraneuronal pathologies is crucial for evaluating potential AD therapeutics^25,26^. A key model in this field involves basal forebrain cholinergic neurons (BFCNs), which are known to be significantly impacted and reduced during the early stages of AD^27^. Current methodologies have enabled the production of age-relevant and AD-specific BFCNs from patient-derived skin cells^28^. However, these cells typically require coculture with primary mouse astrocytes for survival, and the manifestation of Aβ42 and Tau pathologies demands prolonged culture periods, often over 50 days^28^. This study introduces an innovative method that generates BFCNs, fully recapitulating the natural endogenous intraneuronal Aβ42 and Tau pathologies within just 21 days. The early manifestation of these intraneuronal pathologies provides a timely and efficient platform for assessing the efficacy of AD therapeutics, representing a significant advancement in understanding AD treatment strategies.

In this research, we focus on the distinct effects of Adu and LCN on intraneuronal endogenous Aβ42 and pTau. While both antibody drugs target extracellular Aβ42 aggregates, their intraneuronal impacts may differ significantly. This study addresses the urgent need for faster and more effective neuronal models and underscores the importance of intraneuronal pathologies in developing efficacious AD treatments. By exploring the potential disparities in the intraneuronal effects of Adu and LCN, we aim to enhance our understanding of their therapeutic potential and delineate their specific roles in AD treatment, contributing to a more comprehensive approach in combatting this complex disease.

## Results

### BFCN Preparation and Characterization

In our study, basal forebrain cholinergic neurons (BFCNs) were derived from skin cells of both a healthy control (HC, AG08517) and an Alzheimer’s disease (AD) patient (AG06848, Familial PSEN1 mutation), using a commercial BFCN reprogramming kit. This method facilitated a rapid and efficient transdifferentiation of fibroblasts into BFCNs, with over 92% of transduced cells achieving complete conversion within 10 days post-induction (dpi), as detailed in Figure S1. The resulting cells underwent significant morphological changes characteristic of neuronal morphology, including small and round cell bodies with fine neurites (Figure S1). The identity and subtype specificity of the BFCNs were confirmed via antibody staining for general neuronal markers (TUJ1, MAP2, Synapsin 1) and BFCN-specific markers (ChAT, ISL1, TrkA) (Figures 1 and S2). Over 94% of HST+ cells co-expressed MAP2, and more than 95% of MAP2+ cells expressed ChAT and ISL1, indicating a high level of purity and subtype uniformity (Figure 1E and F). The BFCNs also exhibited freeze-thaw viability of 75%–95% and a plating efficiency of 60%–85% within approximately 5 days (Figure S1G). Under our optimized conditions, BFCN monocultures demonstrated robust survival for over 90 dpi, forming very dense networks without the need for coculture with primary mouse astrocytes (Figure S1H).

**Figure 1.**
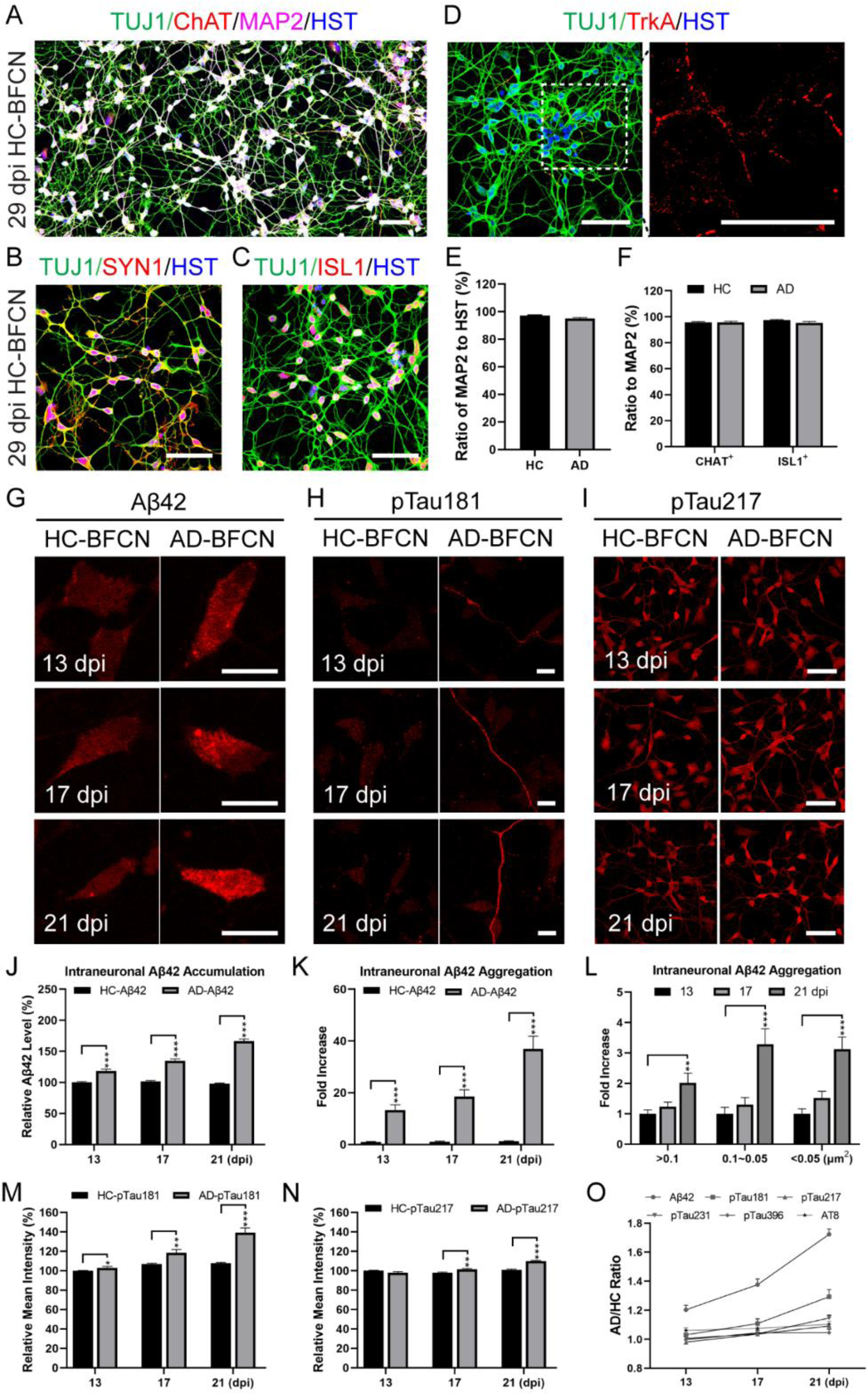
Intracellular Amyloid and Tau Pathology in Neurons Derived from Alzheimer’s Disease Patients. A-D. BFCNs were identified using general neuronal markers (Tuj1, MAP2, Syn1) and BFCN-specific markers (ChAT, ISL1, TrkA). Scale bar, 50 µm. E and F. The BFCNs exhibited high purity (MAP2/HST, HC=97%, n=1154; AD=95%, n=1105) and uniformity (CHAT/MAP2, HC=96%, n=488; AD=96%, n=554; ISL1/MAP2, HC=97%, n=666; AD=95%, n=551) in both healthy controls (HC) and AD patients. G. There was a rapid internal buildup and aggregation of Aβ42 in AD-BFCNs post-conversion and replating. Scale bar, 5 µm. H. An early and progressive increase in the AD biomarker pTau181 was observed in AD-BFCNs, concurrent with Aβ42 accumulation. Scale bar, 20 µm. I. Both HC- and AD-BFCNs showed rapid production and accumulation of pTau217. Scale bar, 50 µm. J. Aβ42 levels in HC- and AD-BFCNs analyzed, normalized to HC at 13 dpi. Mean ± SEM, N=10 random 60× fields (n=105∼117 cells) per time point. K. Fold increase of Aβ42 aggregates in HC- and AD-BFCNs analyzed; normalized to HC at 13 dpi. Mean ± SEM; N=10 random 60× fields (n=222∼298 cells) per point. L. Time-dependent fold increase and particle size of Aβ42 aggregates in AD-BFCNs. Data normalized to particle number at 13 dpi; mean ± SEM; N=10 random 60× fields (n=222∼298 cells) per point. M and N. pTau181 and pTau217 in HC- and AD-BFCNs measured; normalized to HC at 13 dpi. Mean ± SEM; N=10 random 60× fields per point. O. Comparative increase of Aβ42 and pTau forms in AD-BFCNs vs. HC-BFCNs. Mean ± SEM; N=10 random 60× fields per point. Statistical analysis was conducted using a two-tailed unpaired Student’s t-test with GraphPad Prism (Ver. 8) software. Significance levels were indicated as *p<0.05, **p<0.01, and ***p<0.001.

### Pathological Features in BFCNs

To assess the development of pathology in BFCNs, we first demonstrated the specificity of antibodies against Aβ42, various forms of pTau, and the neuronal marker MAP2 (Figure S3). In AD-BFCNs, AD-related pathological features, including the accumulation and aggregation of intraneuronal Aβ42 and the clustering of pTau181^20,21,29^, developed rapidly within 21 dpi (Figure 1G, H, J-M, and O). For aging healthy control BFCNs (66 years), both monomeric and aggregated forms of Aβ42 remained at a low, steady level during culture (Figure 1G, J, and K). In contrast, the burden of Aβ42 in AD-BFCNs (56 years) rapidly increased, reaching 110% at 13 dpi, 120% at 17 dpi, and 160% at 21 dpi compared to HC-BFCNs at 13 dpi (Figure 1J). Simultaneously, the aggregation of Aβ42 within AD-BFCNs drastically accelerated, becoming up to 13, 19, and 37 times higher than in HC-BFCNs at 13 dpi (Figure 1K). Furthermore, the formation of endogenous Aβ42 oligomers and protofibrils in AD-BFCNs, corresponding to small (<0.05 µm²), medium (0.05 – 0.1 µm²), and large (>0.1 µm²) Aβ42 aggregates, exhibited time-dependent patterns and increased sharply, up to threefold by 21 dpi (Figure 1L). Comparative analysis also revealed that pTau181 uniquely accumulated over time in AD-BFCNs (Figure 1H and M), closely correlating with the accumulation and aggregation of intraneuronal Aβ42 (γ=0.999, Figure 1O). Other forms of pTau were present not only in AD-BFCNs but also in HC-BFCNs as early as 13 dpi (Figure 1I and S4). While the progression of other known Tau pathologies, including pTau217, pTau231, pTau396, and pTau202/205 (AT8)^28,29^, was evident in AD-BFCNs, their expression levels in HC-BFCNs also increased from 13 to 21 dpi (Figure 1I, N, O, and S4). Consequently, Aβ42 and pTau181 were chosen as primary indicators for evaluating the effects of LCN and Adu.

### Distinct Effects of LCN and Adu on Intraneuronal Pathologies

We next investigated the intraneuronal impact of LCN and Adu compared to a control immunoglobulin G (IgG, Ctrl) on Aβ42 and pTau181 in AD-BFCNs. Like the control IgG, both LCN and Adu were internalized into the neurons via the Fc gamma receptor Ia (FcRIa) and/or the transferrin receptor (TfR) (Figure 2A and B). Once internalized, LCN localized with intraneuronal Aβ42 aggregates and was partially associated with markers of early endosomes (RAB5), late endosomes (RAB7), lysosomes (LAMP1), and proteasomes (LMP7) (Figure 2C-F). Quantitative immunofluorescence analysis revealed that LCN effectively reduced intracellular Aβ42 levels to about 38% and decreased its aggregates to 70% in AD-BFCNs after 72 hours (Figures 2G-K). Notably, LCN had the most pronounced effect on small aggregates (88%), followed by medium (81%) and large aggregates (58%) (Figure 2K). These results indicate that LCN might selectively bind to the oligomer and protofibril forms of Aβ42 aggregates, forming intracellular antigen/antibody complexes that trigger subsequent degradation. Alongside the reduction in intraneuronal Aβ42 accumulation and aggregation, LCN also significantly reduced pTau181 levels to 20% in AD-BFCNs after a 72-hour treatment (Figures 2H and L). In stark contrast, Adu, which is designed to target fibrillized Aβ42, failed to bind and clear intraneuronal Aβ42 and its aggregates, resulting in no significant effect on the build-up of pTau181 and its axonal accumulation (Figures 2G-L). This lack of efficacy of Adu might explain its limited clinical significance, despite its reported efficiency in reducing extracellular amyloid burden in the brain^9^. Collectively, these findings suggest a potential correlation between the efficacy of these therapeutic antibodies in clearing intraneuronal Aβ42 and their clinical effectiveness in Alzheimer’s disease treatment.

**Figure 2.**
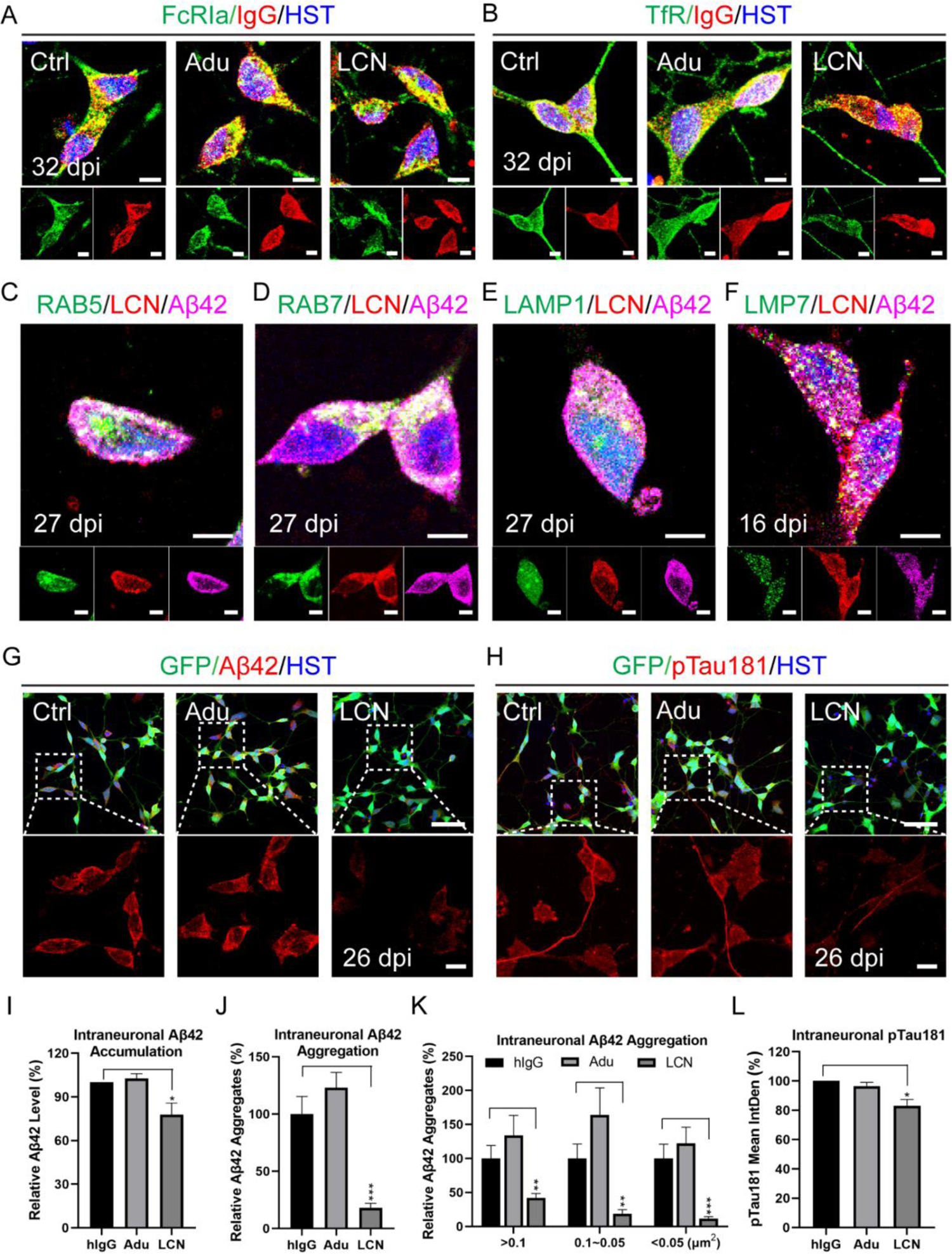
Distinct Impact of LCN and Adu on natural endogenous Aβ42 and pTau181 in AD-BFCNs. A and B. Localization of LCN and Adu in relation to the Fc gamma receptor Ia (FcRIa) and transferrin receptor (TfR) was observed. Scale bar, 5 µm. C-F. LCN was associated with intracellular Aβ42 aggregates and markers of early endosomes (RAB5), late endosomes (RAB7), lysosomes (LAMP1), and proteasomes (LMP7, also known as PSMB8). Scale bar, 5 µm. G. LCN significantly reduced Aβ42 levels and aggregation in AD-BFCNs after 72 hours, a reduction not observed with Adu. Scale bar, 50 µm (upper) or 10 µm (lower). H. LCN was the only treatment that notably decreased pTau181 in AD-BFCNs following 72-hour treatment. Scale bar, 50 µm (upper) or 10 µm (lower). I-K. LCN decreased intraneuronal Aβ42 levels to about 38% and selectively reduced intraneuronal Aβ42 aggregates by 70–90%. Mean ± SEM; N=3 independent samples (Ctrl, n=429; Adu, n=465; LCN, n=434). L. A roughly 20% reduction in pTau181 was achieved with LCN treatment. Mean ± SEM; N=3 independent samples (n=41 random 60× oil fields). Statistical analysis was conducted using a two-tailed unpaired Student’s t-test with GraphPad Prism (Ver. 8) software. Significance levels for LCN versus human IgG (hIgG) were indicated as *p<0.05, **p<0.01, and ***p<0.001.

### LCN/TRIM21-Mediated Clearance of Intraneuronal Aβ42

Tripartite motif-containing protein 21 (TRIM21), a cytosolic IgG Fc receptor with ubiquitin ligase activity, plays a pivotal role in eliminating intracellular antigen/antibody (Ag/Ab) complexes^4^, a crucial mechanism underlying the innovative Trim-Away technology^30^. We hypothesized that TRIM21 might be instrumental in reducing the intraneuronal complexes formed by LCN with Aβ42 aggregates. Notably, after treating AD-BFCNs with LCN, TRIM21 exhibited a slight but significant decline within three days (Figure 3A and B), concurrent with a reduction in Aβ42 (Figure 2I-K). To further explore the degradation pathway of the intraneuronal Aβ42-LCN-TRIM21 complexes, AD-BFCNs were treated with LCN for 72 hours, followed by exposure to either autophagosome-lysosome fusion inhibitor Bafilomycin A1 (BAF) or proteasome inhibitor Bortezomib (BTZ) for an additional six hours. Quantitative immunofluorescence assays revealed a significant increase in both Aβ42 and TRIM21 levels following treatment with these inhibitors compared to the vehicle (Veh) group (Figure 3C-E). These results indicate that both the autophagy-lysosomal and proteasomal pathways contribute to the degradation of the Aβ42-LCN-TRIM21 complexes. Therefore, we conclude that LCN effectively reduces the intraneuronal burden of Aβ42 through the TRIM21 pathway, targeting both autophagic lysosomes and proteasomes.

**Figure 3.**
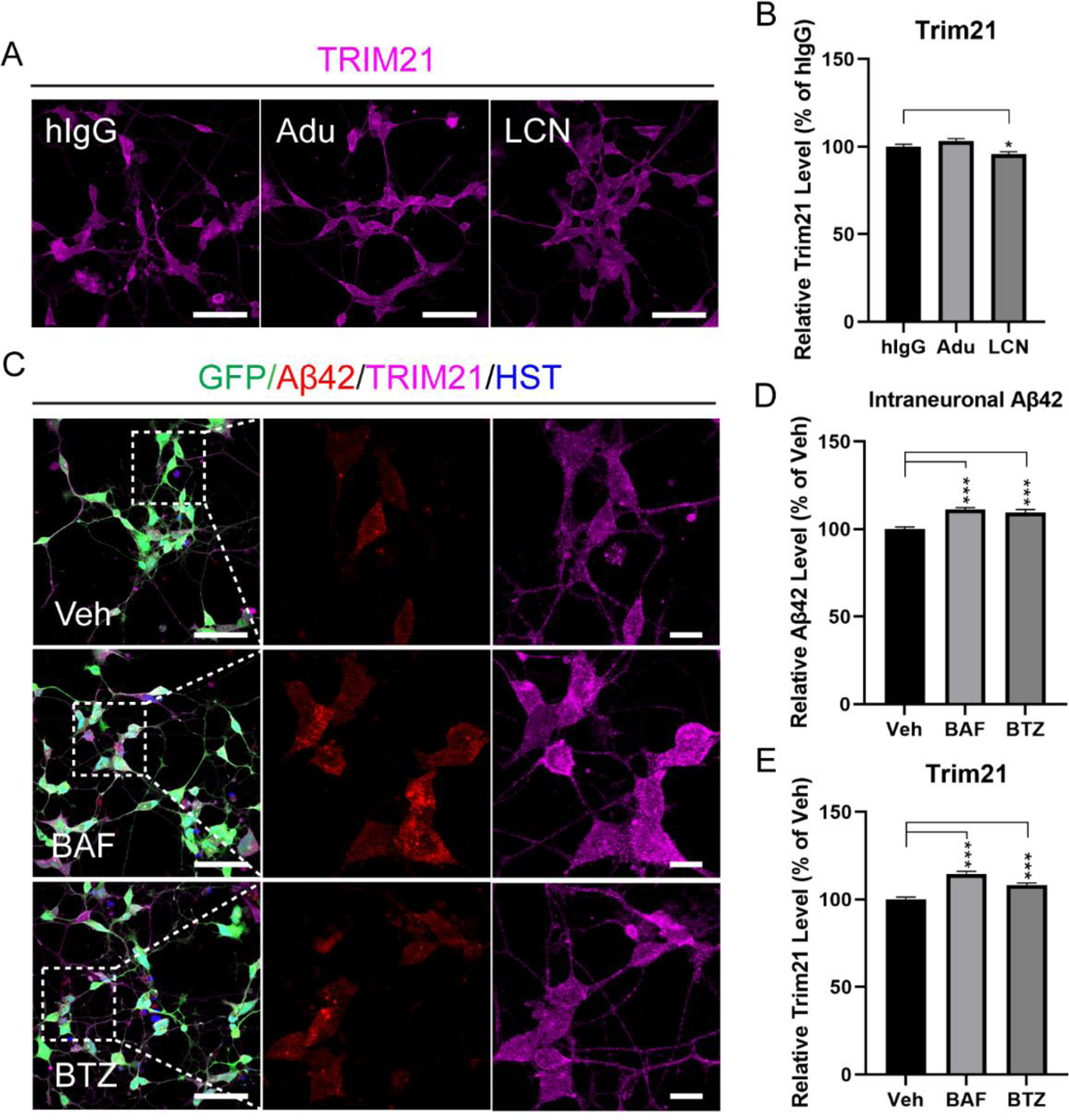
TRIM21-Mediated Clearance of Intraneuronal Aβ42-LCN Complexes. A and B. LCN treatment slightly but significantly reduced TRIM21 levels in AD-BFCNs compared to Adu or control IgG. Mean ± SEM; N=10 random 60× fields. Scale bar, 50 µm. C-E. Following the application of the autolysosome inhibitor BAF (10 nM) or the proteasome inhibitor BTZ (0.5 µM) for 6 hours, both Aβ42 and TRIM21 levels increased in AD-BFCNs. Mean ± SEM; N=10 random 60× oil fields. Scale bars, 50 µm for left panels and 10 µm for right panels. Statistical significance was determined using a two-tailed unpaired Student’s t-test with GraphPad Prism (Version 8) software. Significance is denoted as *p<0.05, **p<0.01, and ***p<0.001.

## Discussion

Our study represents a significant advancement in the development of neuronal models for AD research. By optimizing conversion and culture conditions, we successfully generated BFCNs that rapidly develop Aβ42 and pTau pathologies within 13–21 days, a timeframe considerably shorter than the typical 50-day period^28^. Additionally, these BFCNs were freezable, with freezing/thawing viability comparable to or higher than the iPSC-derived fetal neurons^31^, which significantly facilitates their storage, transportation, and application. This accelerated onset of natural endogenous neuropathology enhances the efficiency and reliability of evaluating potential AD therapeutics, offering a more accessible and dependable platform for testing.

In the context of clinical AD diagnosis, pTau181 and pTau217 are regarded as relatively reliable early pathological biomarkers due to their strong correlation with Aβ42 levels in patients^29,32,33^. In AD-BFCNs, a variety of pathological pTau forms^28,32,33^, including pTau181, pTau202/205 (AT8), pTau217, pTau231, and pTau396, were detected as early as 13 days post-induction (dpi), potentially coinciding with the emergence of Aβ42.

Nevertheless, with the exception of pTau181, similar pTau forms were also observed in HC-BFCNs, even at low Aβ42 levels. Over time, these pathological pTau variants exhibited rapid and significant increases in AD-BFCNs. However, from 13 to 21 dpi, aside from pTau181 which was uniquely present and accumulated in AD-BFCNs, other pTau forms were noticeably present in HC-BFCNs as well. The rate of increase for these pTau forms in HC-BFCNs was notably lower than in AD-BFCNs. While these observations require additional validation across a wider array of samples, they clearly suggest that Aβ42 and pTau181 in AD-BFCNs could reliably serve as indicators for evaluating the efficacy of AD therapeutics.

A core focus of our investigation was the distinct impacts of Adu and LCN on intraneuronal Aβ42 and pTau, shedding new light on their unique roles in combating AD. While both effectively reduce extracellular amyloid-beta in the brain^9–14^, our findings suggest a significant difference in their intraneuronal actions. LCN, designed as a bispecific antibody^14^, may have an increased likelihood of entering neurons compared to Adu. This potential for enhanced neuronal entry facilitates LCN’s formation of cytosolic complexes with endogenous Aβ42 oligomers/protofibrils. It activates the TRIM21 pathway^4,30^, potentially enhancing autolysosome- and proteasome-mediated amyloid clearance, subsequently leading to a considerable decrease in pTau pathology. In contrast, Adu exhibits substantially weaker effects, primarily due to its lower affinity for oligomeric and protofibrillar forms of Aβ42 predominant inside neurons^13^. Because the transferrin receptor is enriched on the neuron membrane^34^, the design of LCN as a bispecific antibody suggests it might not only aid in crossing the blood-brain barrier but also increase its intraneuronal efficacy, thereby possibly boosting its ability to clear neuronal Aβ42.

This disparity underscores LCN’s potential as a more promising therapeutic option, particularly in addressing intraneuronal pathologies closely linked to cognitive decline in AD patients. The limited effectiveness of Adu in reducing intraneuronal Aβ42 and pTau may partially explain its variable clinical results, emphasizing the importance of investigating the intraneuronal impact of potential AD therapeutics.

However, it is crucial to acknowledge the limitations of our proof-of-concept study, which utilized only one cell line from an HC or AD patient. While in vitro models with single or multiple samples provide valuable insights, they may not fully replicate the complex dynamics of AD in vivo. This may explain why we did not observe a significant impact on AD-BFCN viability due to increased Aβ42 and pTau pathologies. In the brain tissue of AD model mice, glial cells activated by pathological Aβ42 and pTau can in turn cause neuronal dysfunction or death^35,36^. Additionally, our investigation does not cover all the underlying mechanisms behind the intraneuronal effects of Adu and LCN, suggesting the need for further exploration.

In conclusion, our research highlights the critical role of intraneuronal pathologies in AD and elucidates the distinct intraneuronal effects of Adu and LCN. These insights emphasize LCN’s potential as a more effective therapeutic agent, paving the way for more comprehensive approaches in AD treatment. The accelerated pathology development in our model offers a crucial platform for future research, underscoring the importance of targeting intraneuronal pathologies in developing effective AD treatments.

## Materials and Methods

### Culture Medium and Reagents

Dulbecco’s Modified Eagle Medium high glucose (DMEM) was sourced from Hyclone (USA). Fetal bovine serum (FBS) was obtained from Corning (USA). Growth factors such as BDNF, GDNF, NT3, and NGF were acquired from Sino Biological (China) or Absin (China). Aducanumab, Lecanemab, autophagy inhibitor Bafilomycin A1, and proteasome inhibitor Bortezomib were purchased from MCE (USA). Control human IgG and forskolin were sourced from Selleck (USA). The BFCN Reprogramming Kit, BFCN Culture Kit, and BFCN Seeding Reagent were supplied by Ecyton Biotech (China). L-glutamine and paraformaldehyde (PFA) were obtained from Aladdin (China). Trypsin, DPBS, DMSO, and other reagents were from Sigma (USA), unless otherwise specified.

### Cell Source and Culture

Human skin fibroblasts, obtained from an aging healthy control (AG08517, 66 years, female) and an AD patient (AG06848, 56 years, female, Familial PSEN1 mutation), were sourced from the Coriell Cell Repositories (USA). These fibroblasts were maintained in fibroblast medium (FM), which is high glucose DMEM enriched with 15% fetal bovine serum (FBS), until they reached confluency. Subsequently, they were trypsinized for splitting and lentiviral transduction.

### Transdifferentiation

Direct conversion was carried out using the protocol provided in the BFCN Reprogramming Kit (ECT-BFCNR01). Upon reaching confluency, fibroblasts were split and plated at 1.5 × 10^4^ cells per cm^2^ in vessels pre-coated with coating buffer (ECT-CB). After overnight culture in Fibroblast Medium (FM), fibroblasts were exposed to lentiviruses (LV-BFCN-TFs) carrying key transcription factors such as NKX2.1 and LHX8 for 24 to 48 hours. Post-infection, cells were placed in fresh FM for a day to recover from viral stress. Conversion began when cells were switched to Neuronal Induction Medium (ECT-NIM) containing NIM supplement (ECT-NIMS) and 1× Golden Solution (ECT-GS), marking 0-day post-induction (dpi). GS was formulated with a cocktail of key reprogramming chemicals and reagents confidential to Ecyton Biotech, which ensures high conversion efficiency. The complete NIM was half-refreshed at 2, 4, and 6 dpi. Live cell imaging and monitoring of the conversion process were performed using a Leica DMIL Inverted Fluorescence Microscope. Conversion efficiency was assessed by the percentage of TUJ1+ neurons among GFP+ cells at 10 dpi.

### BFCN Purification, Culture, Cryopreservation, and Recovery

Converted BFCNs were purified between 8 to 10 dpi using established protocols^37^. It is important to note that both high transduction rate and conversion efficiency are crucial for achieving both high purity and substantial yield through this protocol. For culturing, the optimized BFCN Culture Kit (ECT-BFCNCM01) was utilized. BFCNs were plated at a density of 10 × 10^4^ cells per cm^2^ on glass coverslips or in 24-well plates pre-coated with the neuronal coating reagent (ECT-CR), and then cultured in neuronal medium (ECT-NM) supplemented with 1 mM L-glutamine, 5 µM forskolin, 30 ng/ml NGF, and 10 ng/ml each of BDNF, GDNF, and NT3. The medium was half-renewed every 4-6 days until ready for assessment. Following purification and culture on coverslips for the specified durations, neuronal purity was evaluated by the percentage of MAP2+ neurons among HST+ cells. Subtype homogeneity was determined by the percentage of ChAT+ or ISL1+ neurons among MAP2+ cells.

For cryopreservation, BFCNs were centrifuged at 500 g for 5 minutes and resuspended in 50 µl NM per 100 × 10^4^ cells. After prechilled at 4℃ for 5 min, the cell suspension was mixed with 200 µl cold CryoStor® CS10 medium (STEMCELL Technologies, Canada) and then transferred into cryovials (Nalgene, USA) or substitutes. This suspension was kept at 4℃ for 10 minutes before being frozen at −80℃ in a BeyoCool™ container (Beyotime, China) and ultimately stored in the vapor phase of a liquid nitrogen tank for long-term preservation.

For recovery, BFCNs were revived using the BFCN Culture Kit (ECT-BFCNCM01) and Seeding Reagent (ECT-SR), which are part of a proprietary culture and seeding system developed by Ecyton Biotech (China) to enhance cell viability and growth. This system comprises a specially formulated mix of small molecules and reagents, optimized for the efficient recovery of BFCNs. Culture vessels were pre-coated with ECT-CR overnight.

The cryovial was carefully and quickly thawed in a 37℃ water bath with gently swirling and then placed on ice or in a cold box immediately before the last piece of ice disappeared. The complete NM described above, further enriched with SR, was added to dilute the suspension approximately 5-fold. Thawing viability was immediately checked with Trypan Blue (Sigma, USA). Cells were plated on coated vessels and incubated at 37℃ for 15-30 minutes. After removing the seeding medium containing DMSO, fresh NM with SR was added gently. Cells were maintained for 5 days, then continued in regular NM without SR as described. Plating efficiency was evaluated at the specified time point using the live cell imaging technique described above.

### Treatment with Monoclonal Antibodies and Chemicals

AD-BFCNs were grown on coverslips in 24-well plates, each well containing approximately 1 ml of culture medium, until the cells essentially formed a neural network. Monoclonal antibodies, including control human IgG, Adu, and LCN, were diluted to 2 µg/ml in the medium before being introduced to the cells to reach a working concentration of 1 µg/ml. Following gentle agitation, the cultures were returned to the incubator to continue growing for 72 more hours. Subsequently, for chemical intervention, cells pre-treated with antibodies for 72 hours underwent further treatment with autophagy and proteasome inhibitors—Bafilomycin A1 (BAF, 10 nM) and Bortezomib (BTZ, 0.5 µM) respectively—diluted to their intended final concentrations in the culture medium, utilizing DMSO as a vehicle control (Veh). After another round of gentle mixing, the cultures were incubated for an additional 6 hours. Finally, cells were fixed using 4% PFA to ready them for immunocytochemical staining and further assessments.

### Immunocytochemistry

Cells fixed with 4% PFA for 15 min at room temperature (RT) and rinsed twice with PBS buffer were subjected to immunocytochemistry according to previously described protocols^37^, with slight modifications. Briefly, fixed cells were permeabilized and blocked in PBS containing 3% BSA and 0.2% Triton X-100 for 2 hours at RT. Next, primary antibodies diluted in the blocking buffer were applied for 2 hours at RT, followed by three 10-minute washes with PBS buffer containing 0.2% Triton X-100. Subsequently, secondary antibodies were applied for 1 hour at RT, followed by three 10-minute washes with the wash buffer. Details regarding the source, supplier, dilution, and other pertinent information about the primary and secondary antibodies used in this study are provided in Table S1. Nuclei were counterstained with Hoechst 33342 (APExBIO, USA). Coverslips were then mounted on glass slides using anti-fade fluorescence mounting medium.

### Image Capture and Processing

High-quality images from random 60× oil immersion fields were captured with an Olympus FV3000 confocal microscope using optimal and consistent settings for each channel. Z-scanning was performed over a 4.2 µm range with 0.6 µm intervals between steps. The initial processing of Z-scanned images utilized the microscope vendor’s software. Subsequent processing and analysis were carried out with ImageJ software. For visualization of Adu and LCN internalization pathways in BFCNs, and their interaction with markers of early endosomes (RAB5), late endosomes (RAB7), lysosomes (LAMP1), and proteasomes (LMP7, also known as PSMB8), a single Z-step that provided the clearest signal was selected. This step was then presented with an optimal threshold for clarity.

### Quantification of Immunofluorescence Intensity

To ensure consistent analysis, images from the same experimental set were identically adjusted using ImageJ. For quantifying intraneuronal Aβ42 accumulation, projected images from 2 to 5 Z-steps were directly analyzed. The soma boundary of each cell was primarily delineated using the magic wand tool, with some manual tracing, after which both mean and total fluorescence intensities were measured with the analyze tool. For Aβ42 aggregate analysis, each projected image was converted to 8-bit format and thresholds were set to an optimal range. Aggregate counts were obtained using the analyze particles tool and data were exported to Excel for classification into three size categories based on their area: small (<0.05 µm²), medium (0.05-0.1 µm²), and large (>0.1 µm²).

For quantification of intraneuronal pTau pathology variations, all 8 Z-step encompassed projected images were analyzed with customized threshold adjustments for each pTau pathology. The analyze tool was employed to measure both mean and total fluorescence intensities for all cells in each image. Intracellular TRIM21 level quantification was performed using similar methods applied for pTau analysis.

### Statistical Analysis

Each experiment was independently replicated a minimum of three times. Data are presented as mean ± standard error of the mean (SEM), with the total number of experiments (N) and the number of observations or cells within those experiments (n) specified. Statistical analyses were performed using a two-tailed unpaired Student’s t-test via GraphPad Prism (Version 8) software. Levels of significance are denoted as follows: *p<0.05, **p<0.01, and ***p<0.001.

## Author Contributions

Conceptualization, M.-J.L., Z.L., and M.-L.L.; Methodology, M.-J.L., Z.L., F.L., X.S., Y.-F. S., and M.-L.L.; Investigation, M.-J.L., Z.L., F.L., and X.S.; Data analysis and interpretation, M.-J.L., Z.L., F.L., X.S., Y.-F. S., and M.-L.L.; Resources, M.-J.L., Y.-F. S., and M.-L.L.; Writing – draft, M.-J.L. and Z.L.; Writing – Review & Editing, M.-L.L.; Supervision, M.-J.L. and M.-L.L. All authors reviewed and approved the final manuscript.

## Acknowledgments

We thank the Coriell Cell Repositories for providing human fibroblasts. Additionally, we extend our gratitude to ChatGPT for the English language editing of our manuscript.

---

**Figure S1.**
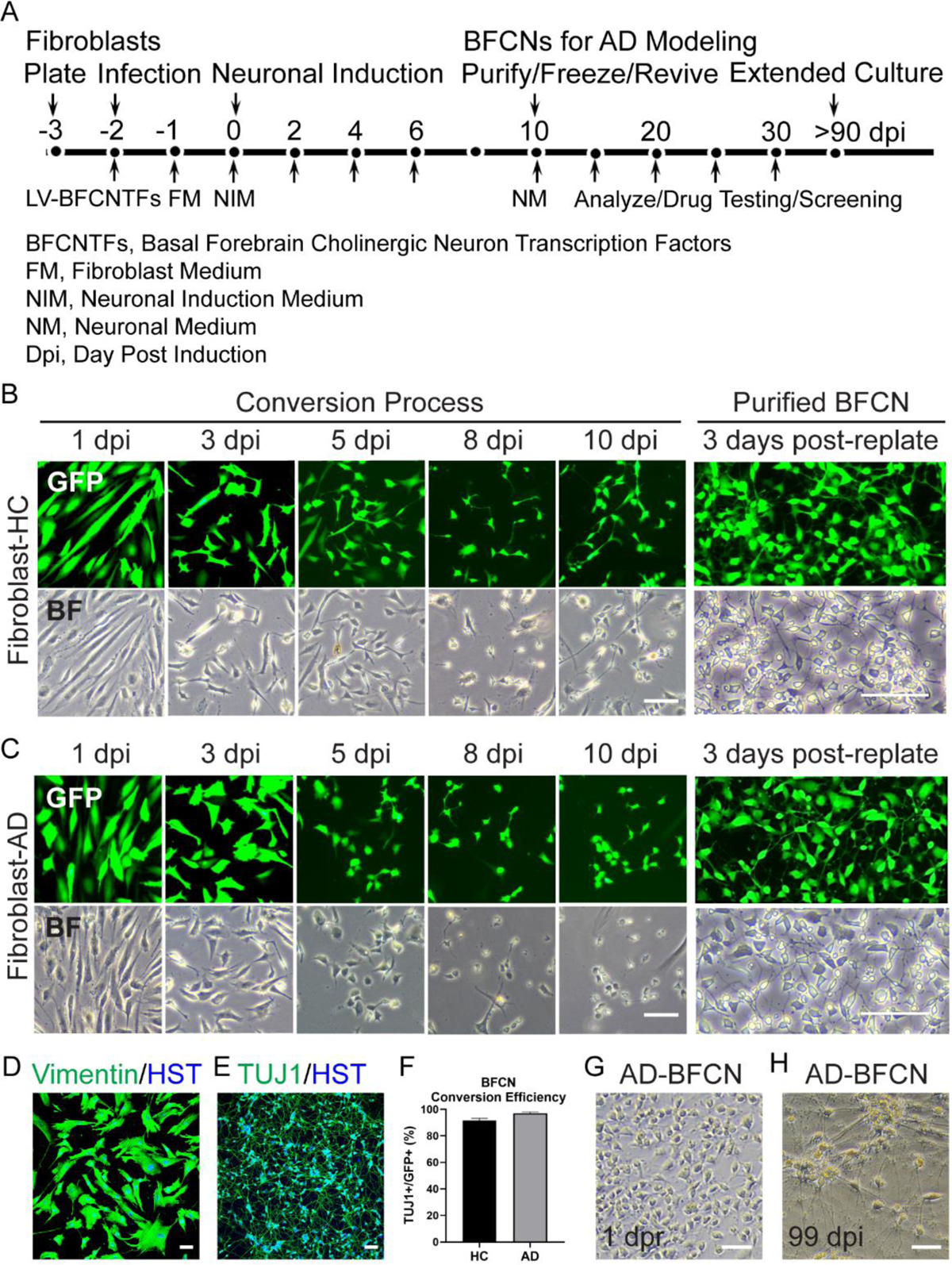
Rapid Conversion of Fibroblasts from Healthy Controls (HC) and AD Patients into Induced BFCNs. A. The direct conversion process from patient fibroblasts to BFCNs within 10 days post-induction (dpi) is represented schematically. B and C. Morphological changes are illustrated during the stages of neuronal conversion and purification. Scale bar, 100 µm. D and E. Comparative images display the original vimentin-positive fibroblasts and the resultant highly pure TUJ1-positive BFCNs. Scale bar, 50 µm. F. High conversion efficiency was observed in both HC (92%) and AD (96%) fibroblast samples. Quantitative data are presented as mean ± standard error of the mean (SEM), based on N=10 random 20× fields (HC, n=436; AD, n=273). G. The BFCNs showed the ability to be frozen and subsequently thawed with minimal loss of viability. ‘Dpr’ refers to days post recovery. Scale bar, 50 µm. H. In monoculture, the BFCNs exhibited robust survival for over 90 days post-induction (dpi), forming very dense neuronal networks. Scale bar, 50 µm.

**Figure S2.**
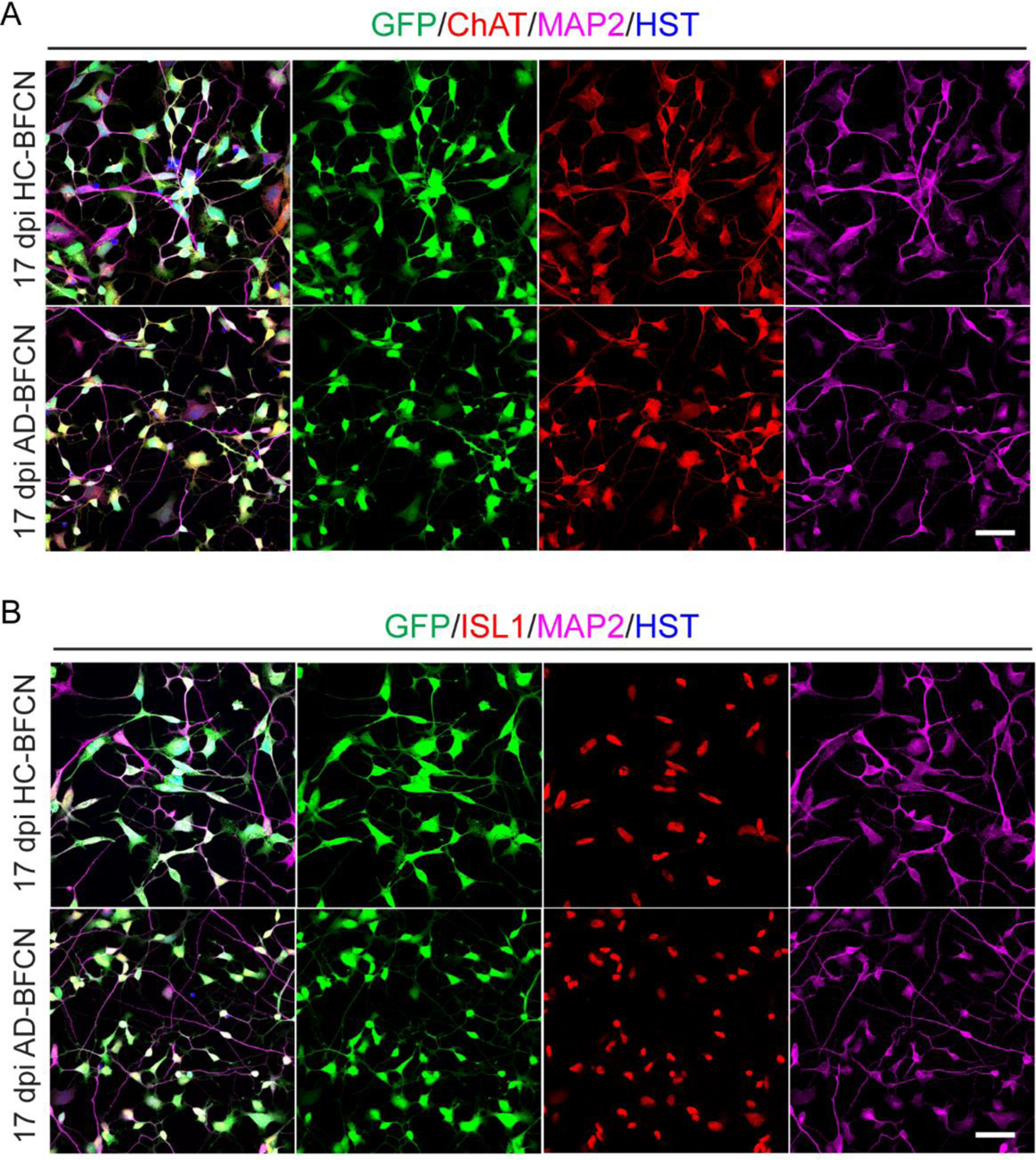
High Purity of BFCNs Derived from Healthy Control (HC) and AD Patient Fibroblasts. A and B. These panels show representative images of highly pure BFCNs derived from HC and AD patient fibroblasts. The images demonstrate the co-expression of the mature neuronal marker MAP2 with cholinergic neuronal markers ChAT and ISL1 in both HC- and AD-BFCNs. Scale bar, 50 µm.

**Figure S3.**
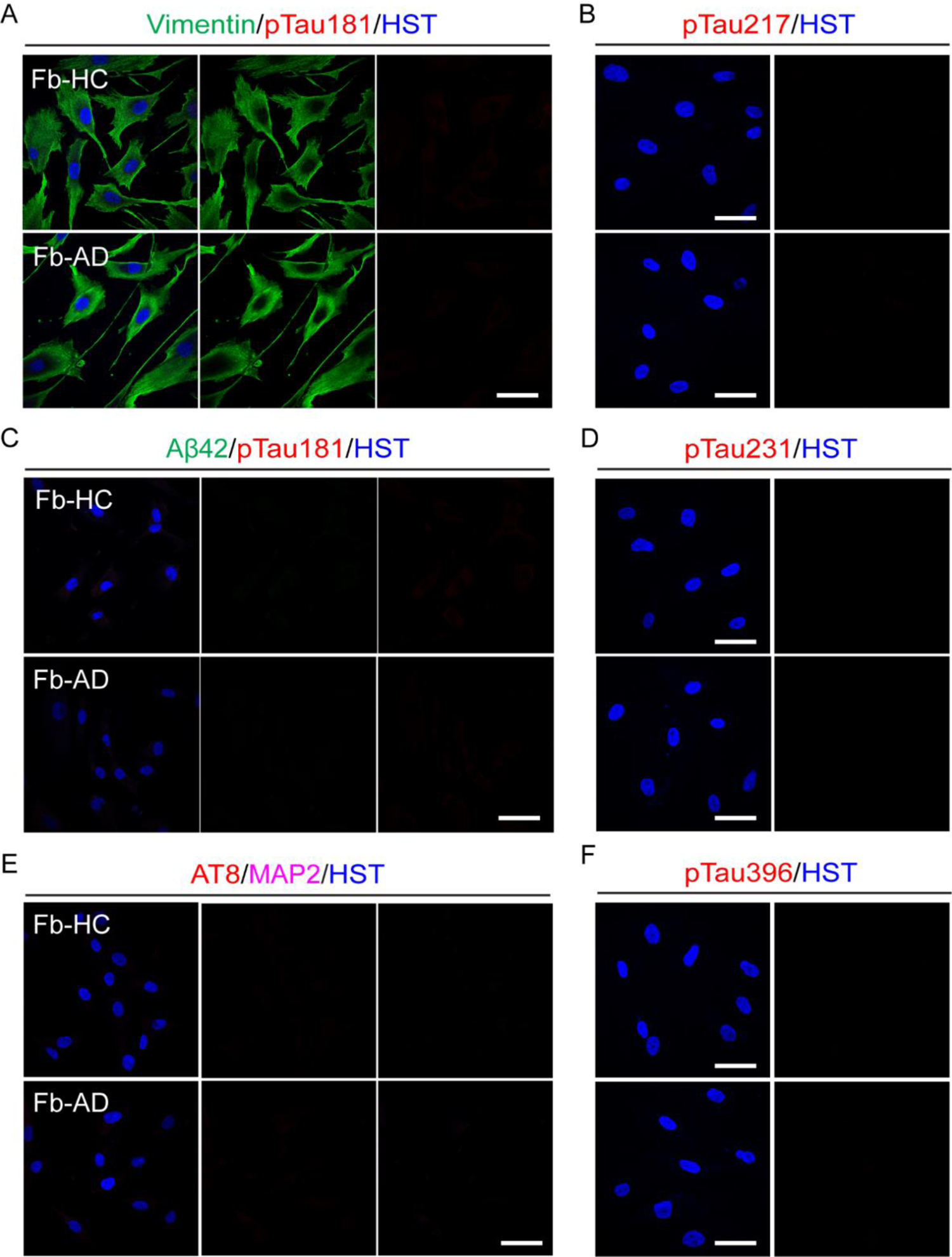
Specificity Confirmation of Antibodies Against Aβ42, Various Forms of pTau, and the Neuronal Marker MAP2. A-F. These panels illustrate the absence of positive staining for Aβ42, pTau181, pTau217, pTau396, AT8, and MAP2 in vimentin-positive fibroblasts derived from both a Healthy Control (HC) and an Alzheimer’s Disease (AD) patient. This lack of staining in fibroblasts, which do not express these neuropathological or neuronal markers, confirms the specificity of the antibodies used. Scale bar, 50 µm.

**Figure S4.**
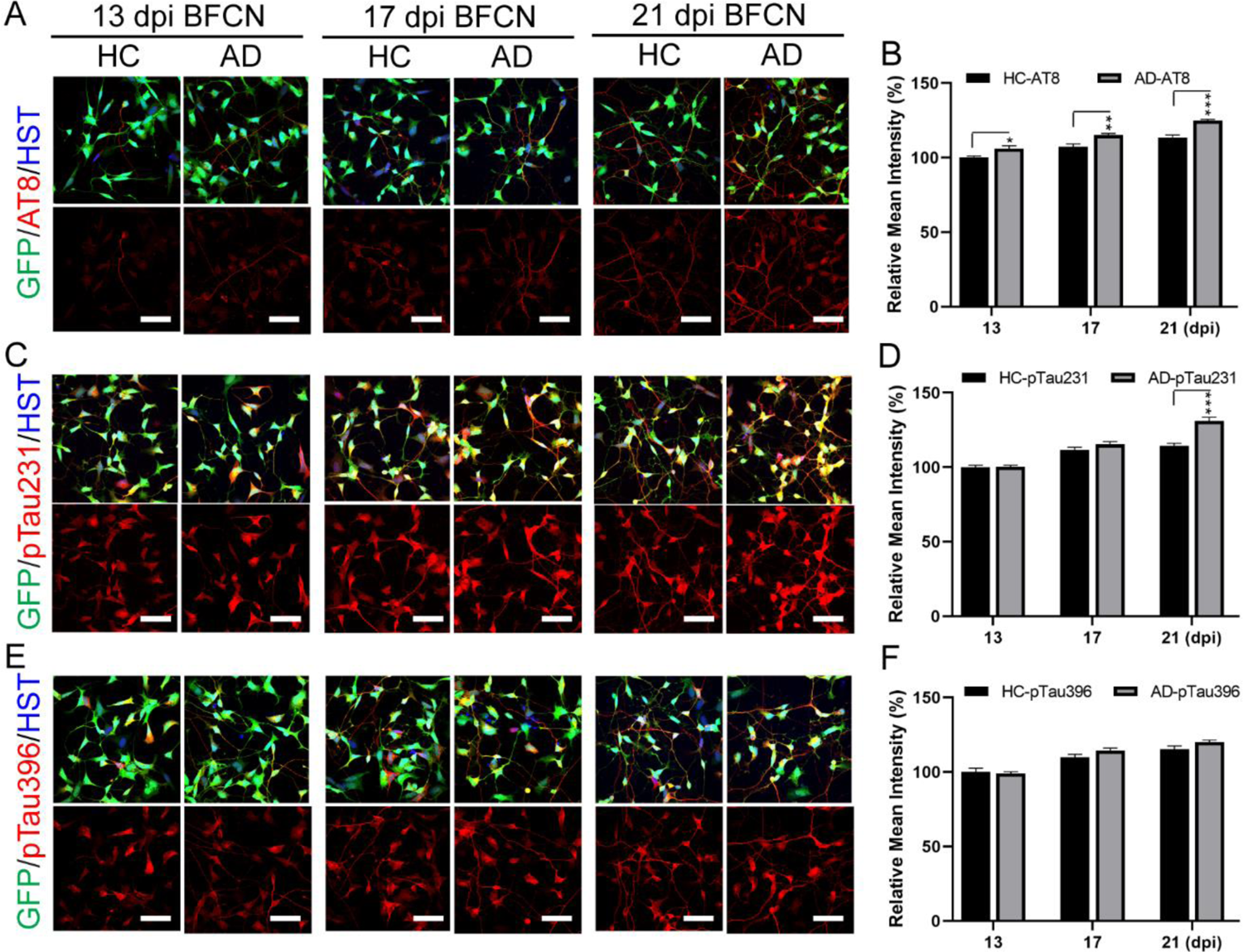
Varied Tau Pathogenesis in Healthy Control (HC) and AD Patient-Derived BFCNs. A and B. These panels show a gradual increase in AT8 staining in both HC- and AD-BFCNs, with a more pronounced increase observed in AD-BFCNs compared to HC-BFCNs from 13 to 21 days post-induction (dpi). Scale bar, 50 µm. C and D. Both HC- and AD-BFCNs exhibit strong and increasing expression of pTau231 from 13 to 21 dpi. Notably, the level of pTau231 was significantly higher in AD-BFCNs than in HC-BFCNs at 21 dpi. Scale bar, 50 µm. E and F. Identical and gradually increasing patterns of pTau396 expression were observed in both HC- and AD-BFCNs from 13 to 21 dpi. Scale bar, 50 µm. Quantitative data were normalized to the HC group at 13 dpi and are presented as mean ± SEM. N=10 random 60× oil fields. Statistical significance was determined using a two-tailed unpaired Student’s t-test with GraphPad Prism (Ver. 8) software. AD versus HC, *P<0.05, **P<0.01, ***P<0.001.

**Table S1.** Primary and secondary antibodies used in this study.

| **Primary Antibody** | **Host** | **Identifier** | **Supplier** | **Dilution** |
| --- | --- | --- | --- | --- |
| pTau181 | Mouse | 846602 | Biolegand, USA | 1:2000 |
| Aβ42 | Rabbit | 24090S | CST, USA | 1:2000 |
| Aβ42 | Mouse | 8005509 | Biolegand, USA | 1:2000 |
| Synapsin1 (SYN1) | Rabbit | D12G5 | CST, USA | 1:500 |
| CHAT | Rabbit | A5303 | Bimake, USA | 1:500 |
| MAP2 | Chicken | Ab5392 | Abcam, England | 1:5000 |
| TUJ1 | Mouse | 66375-1-1g | Proteintech, USA | 1:10,000 |
| TRKA | Rabbit | 11073-R024 | Sino Biological, China | 1:500 |
| ISL1 | Rabbit | A5647 | Bimake, USA | 1:2000 |
| Vimentin | Rabbit | A5862 | Bimake, USA | 1:10,000 |
| FCGR1A (FcRIa) | Rabbit | A02428-3 | Boster, USA | 1:500 |
| Transferrin Receptor | Rabbit | PB9233 | Boster, USA | 1:500 |
| Rab5 | Rabbit | A5992 | Bimake, USA | 1:1000 |
| Rab7 | Rabbit | A5661 | Bimake, USA | 1:1000 |
| LAMP1 | Rat | sc-19992 | SANTA CRUZ, USA | 1:250 |
| pTau202/205 (AT8) | Mouse | MN1020 | Invitrogen, USA | 1:2000 |
| pTau217 | Rabbit | A00896-40 | GenScript, USA | 1:500 |
| pTau231 | Rabbit | A5950 | Bimake, USA | 1:2000 |
| pTau396 | Rabbit | A5952 | Bimake, USA | 1:2000 |
| LMP7 (PSMB8) | Mouse | AG2982 | Beyotime, China | 1:200 |
| TRIM21 | Rabbit | 12108-1-AP | Proteintech, USA | 1:1000 |
| **Secondary Antibody** |  |  |  |  |
| Anti-Mouse IgG.488 | Donkey | 715-545-150 | Jackson, USA | 1:500 |
| Anti-Mouse IgG.647 | Donkey | 715-605-150 | Jackson, USA | 1:500 |
| Anti-Rabbit IgG.594 | Donkey | 711-585-152 | Jackson, USA | 1:500 |
| Anti-Human IgG.594 | Goat | 109-585-003 | Jackson, USA | 1:500 |
| Anti-Chicken IgY.488 | Donkey | 703-545-155 | Jackson, USA | 1:500 |
| Anti-Chicken IgY.647 | Donkey | 703-605-155 | Jackson, USA | 1:500 |
| Anti-Rat IgG.488 | Donkey | 712-545-150 | Jackson, USA | 1:500 |

